# Murine infection by *Mycobacterium marinum* is a reliable model for Bone and Soft-Tissue Damage

**DOI:** 10.1101/2022.08.18.504251

**Authors:** Mahendra Kumar, Ramaraju Ambati, Prachi J Urade, Anil Lotke, Musti Krishnasastry

## Abstract

Extra-pulmonary tuberculosis (EPTB) constitutes 15-20% of the entire TB cases worldwide, and immune-suppressive conditions like HIV-AIDS further aggravate the disease often without symptoms and lack of proper diagnostic method delays the treatment. A thorough understanding of the EPTB infection and the pathogenesis is necessary and requires a reliable in-vivo animal model that mimics pathology similar to human infection. The *M. marinum* mice infection model presented here offers visible and quantifiable pathological features. Moreover, sections of the infected tails exhibited infiltration of the immune cells, a prominent feature frequently observed. Interestingly, the micro-CT imaging of the infected mice’s tails displayed bone erosion to the extent of the coccygeal vertebral loss. Furthermore, infection of the mice with drug-resistant such as Isoniazid (IRP) and Ethambutol (EmbRP) of *M. marinum* populations exhibited pathological features akin to wild-type *M. marinum* infection. At the same time, for EmbRP, the severity is significantly reduced, suggesting the nature of the selected population and its ability to retain or fix the virulent determinant(s) during bacterial growth. These findings advocate the use of the developed model to understand the EPTB precisely bone and spine TB, and it can be further utilized to develop novel therapeutics and diagnostics.

## Introduction

*Mycobacterium tuberculosis*, undoubtedly, is an evolutionarily adapted organism causing the human tuberculosis disease [1]. *M. tuberculosis* is a member of the mycobacterium complex (MTBC) that includes mycobacteria capable of causing tuberculosis like disease in their respective host; like bovine pathogen *M. bovis*, rodent pathogen *M. microti* and human pathogens like *M. africanum*, and *M. canettii* [2–7]. The members of MTBC complex shares close homology at the genetic level and the manifestation of the disease symptoms to their respective hosts are very much identical. Most of the MTBC members infect humans as the case of zoonosis but they can’t establish the primary niche in the lungs but leads to extra-pulmonary tuberculosis, e.g. *M. bovis* infects and colonizes in the gut lumen of the humans and manifests the pathology; while *M. microti* colonizes in lymph nodes, peritoneal layer together with lungs [3,8–10].

The spread of *M. tuberculosis* is mainly through aerosol association that limits the use of it in regular laboratory practices. The current regulations require higher level of containment facility such as bio-safety level-3 (BSL-3) environment that requires capital intensive investment, expertise and training [11–13]. Investigators have often chosen hosts such as BCG, *M. smegmatis* to gain deeper understanding of the onset of the tuberculosis disease by expression of molecules/modules in them [14,15]. Although, use of these hosts has enriched our understanding of the role of numerous mycobacterial genes in the disease manifestation but it is limited to the study of the role of individual gene(s), at best, as such an approach cannot provide all the mechanisms that operate in tandem. Similarly, attenuated *M. tuberculosis H37Ra* was often used to understand the role of different genes in the virulence and its main utility is to address the mechanism involved in its survival inside the host as it can survive for a long periods of time without any visible symptoms of virulence [8,16,17]. Although, *M. tuberculosis; H37Ra* and *H37Rv* are genetically related and thought to have originated form same parental strain, the use of H37Ra as a model organism is however, limited especially, where, the involvement of the endogenous machinery in establishment of disease is concerned as many pathological features are not similar to that elicited by H37Rv [18–20].

*Mycobacterium marinum*, like the human pathogenic agent, is also an intracellular, acid fast, non-tuberculous-mycobacteria that causes tuberculosis-like disease in poikilotherms, primarily in fish; it can manifest clinical symptoms in frogs, eels, oysters, toads, and snakes [21,22]. The recommended growth temperature is ∼30°C, as per American Type Culture Collection while it was also shown to grow at 35°C and above, with in-vitro doubling time ∼4 hours. Phylogenetically, *M. marinum* exhibits 99.3% 16SrRNA sequence homology with *M. tuberculosis* [23–26]. The phylogenetic closeness of *M. marinum* is very well translated to its etiology as it also infects host macrophages, forms caseating granuloma, similar to that formed by the *M. Tuberculosis* [22,27,28]. Interestingly, the virulence determinants of both *M. tuberculosis* and *M. marinum* are easily swappable; suggesting that most of the mechanisms that operate or regulate the virulent determinants might be conserved between these strains [22,29]. Furthermore, it occasionally infects the extremities of the swimmers, swimming pool workers (non-chlorinated), personals involved in the aquaculture or aquaria, residents of cold surroundings and immune-compromised individuals like HIV-AIDS patients. *The M. marinum* causes nodules and skin lesions termed as nodular granulomatous or skin granulomatous disease that is usually limited to skin but sometime it may exhibit the systemic infection [26,30].

In comparison to other strains discussed above, *M. marinum* is relatively safe, easy to propagate in laboratory setup and requires BSL2 level. Moreover, its faster growth rate and ability to manifest symptoms in the gut and visceral organs like liver, kidney or spleen including musculoskeletal deformities that leads to spinal curvature in zebra fish. These properties further enhances its use as a model organism to understand the progression of tuberculosis like disease in fish and amphibians in regular laboratory setup by using zebrafish as the animal model [22]. The infection of mice with *M. marinum* is relatively easy by following well documented animal handling practices. Many pathological observations on soft tissue damage, bone erosion of the coccygeal vertebrae and formation of the caseating granuloma can easily be correlated to the human infection owing to the extendibility of mice background to humans [31].

The property of *M. marinum* to cause skin lesions in humans, and bone erosion to the mice is further utilized to reinforce earlier observations to understand the extent of applicability of mouse model infection by *M. marinum*. This approach can be used to monitor the progression of skin granulomatous lesions to bone and spine TB in humans and at the same time such a model will strengthen the understanding regarding strategies employed by *mycobacterium species* over the course of zoonosis and *M. tuberculosis* infection to humans.

## Materials and methods

All mice experiments were carried as per protocols approved by Institutional Animal Ethical Committee, National Centre for Cell Science, Pune.

### Bacterial culture and antibiotic selection

*M. marinum, H37Ra* and *M. smegmatis* were grown in 7H9 broth base medium supplemented with 10% ADC (Albumin fraction-V, Dextrose and catalase) and 0.05% Tween-80 to mid-logarithmic phase (A_600_∼0.3-0.4). The cells were harvested by centrifugation at 3500g for 10 minutes at 4°C. *M. marinum* was also cultured in the presence of antibiotics Ethambutol (EMB) and Isoniazid (INH); briefly 10 μl (bacteria equivalent to 5×10^5^ CFU) was inoculated from the glycerol stock in 25 ml 7H9 medium and grown to mid-logarithmic phase (A_600_∼0.45-0.55). The resultant culture was passaged from lower to higher concentration of antibiotics through gradual increase i.e. concentration ranging from sub-lethal (0.05 μg/ml,) to above MIC_90_ (8 μg/ml in case of EMB and 200 μg/m for INH). The resultant populations at each stage were preserved and the highest concentration were used for the experiments described here and are termed as EmbRP and IRP. The EmbRP was further grown after withdrawal of EMB to ascertain the growth kinetics. This population is termed as the EmbRP-Rev.

### Mice infection

C57BL/6 female mice of 6-8 weeks age were infected with 100μl bacterial suspension in phosphate buffered saline (PBS), equivalent to 1×10^7^ bacterial cells via tail vein injection. For infection, harvested cells were washed thrice with PBS by centrifuging it at 3500g for 10 minutes at 4°C. The bacterial suspension was adjusted to A_600_∼1.0-1.2 carrying cells equivalent to 1×10^8^ in 1ml PBS which was used for infection.

### Analysis of tail lesions and bone erosion

The infected mice tails were subjected to micro-computed tomography (Micro-CT) scanning with the help of SkyScan 1276 image scanner; Bruker-MicroCT, Kontich, Belgium. With the scanning parameters adjusted to pixel size 20 μm with 50 kV source voltage and 100 μA source current with the 0.5 mm Al filter. The obtained images were further reconstructed by using NRecon. Whereas, CTAn and CTVol were employed to analyze the reconstructed images and reconstructing the 3D models of the tail sections respectively.

### Histological analysis of the tail lesions

The infected mice tails and lungs were preserved in the 10% formalin. The preserved mice tails were de-calcified, followed by the preparation of the paraffin blocks of the de-calcified tails and the preserved lungs to obtain the sagittal sections of 0.5μm thickness across the lesion length. The obtained sagittal sections were stained with Hematoxylin and Eosin for the histological analysis.

The sections were visualized under the light microscope with 10x objective and microphotograph was recorded using Nikon microscope.

## Results and discussion

### *M. marinum* infection to mice

Infection of mice tails with *M. marinum* causes multiple tail lesions of different length across the tail length not exclusively confined to the site of the infection. The lesions are characterized by the pus filled regions. The visible appearance of the lesion on the mice begins on 4-7 days with the development of the edema which subsequently develops into the lesions (Fig. 1A). The lesions seen by us appear to be very similar to earlier observations by Eric Brown’s group [31]. In the extreme cases of infection, tail deformation is frequently observed. The deformation of the tail is mainly associated with the excessive inflammation and pus filled lesions on either side of the tail as evident through visual and histopathological observations. Furthermore, the tail tip or a region of the tail dissociates from the body due to severe necrosis and it can happen anywhere in the tail. The severity of the symptoms enhances with the onset of the infection and peaks at around 15-18 days. The mice tails infected with *M. smegmatis mc*^*2*^ *155* and *H37Ra* displayed typical appearance with no lesion formation at all (Fig 1B and 1C). The absence of any noticeable pathological features to the *M. smegmatis mc*^*2*^ *155* and *H37Ra* infected mice tails suggests the inability of these mycobacteria to efficiently colonize at the tail owing to their attenuated and/or non-pathogenic nature. Moreover, *H37Ra* may exhibit the selective localization to other parts of the body such as lungs, where the environment is more conducive to its growth/survival. The emergence of tail lesions in *M. marinum* infected mice tails advocate its virulent nature, and also explains its ability to cause skin nodules to the individual living in the temperate climate. The localization of the lesion exclusively to the tails and not to other organs is best explained by lower temperature of the tail which is generally lower than that of other organs. The observed pathological features of the mice tails are graded on a pathological score from 1-4 where, maximum severity is set at 4. To grade the observed phenotypes different parameters are considered like the nature of lesion, length of lesion and the morphology of the tail as described in the Table 1, and plotted on a pathological score (Fig. 1D).

**Fig. 1:**
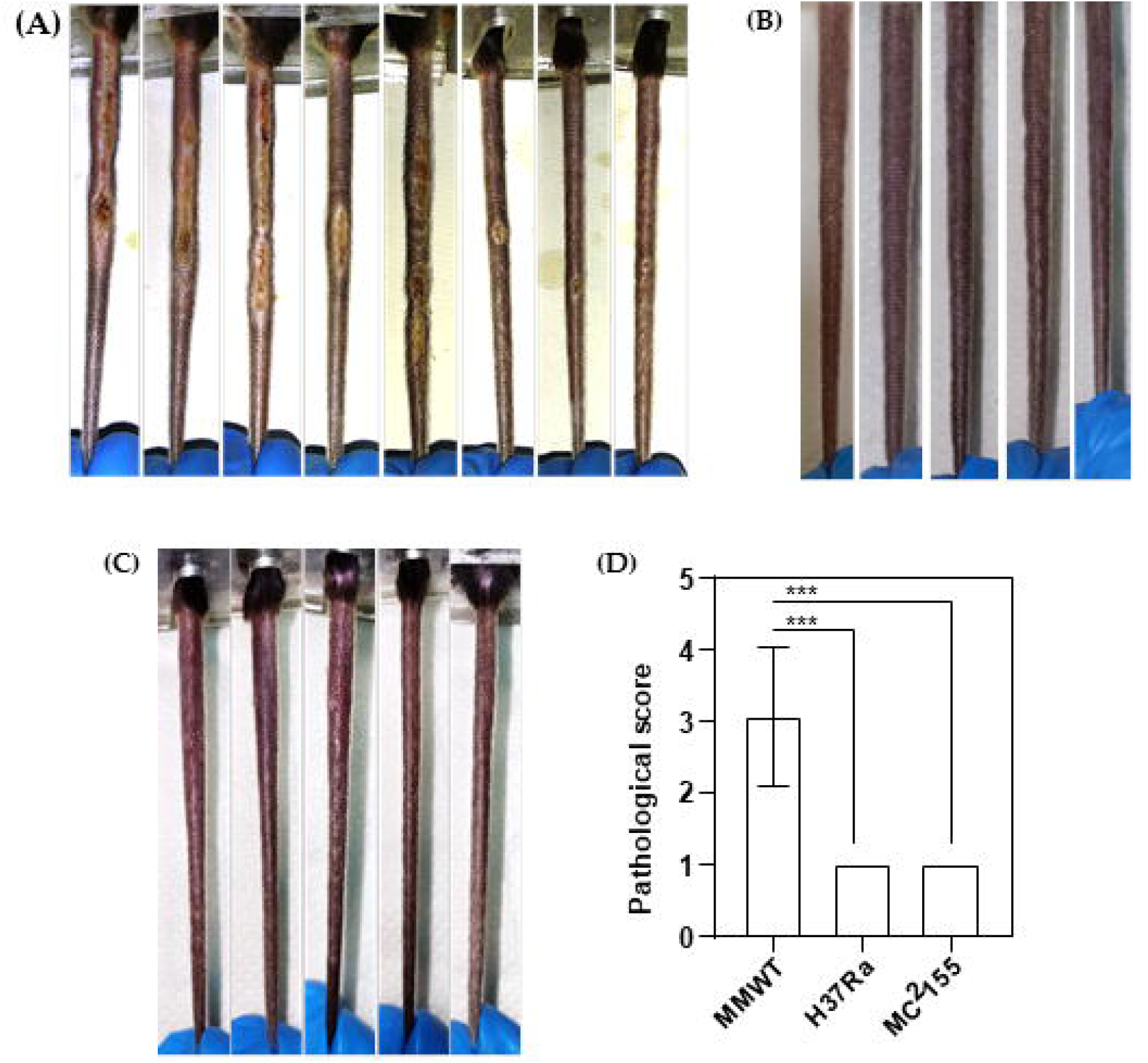
Infection of Mice tails with mycobacterial strains. (A): *M. marinum*, (B): *H37Ra* and (C): *M. smegmatis mc*^*2*^ *155*: C57BL/6 mice were infected with 1*10^8^ bacterial cells via tail-vein injection. Mice were partially restrained using standard restrainer and the injections were performed as described in methods section. (D): The panel D represents quantitation of the severity of lesions of the infected mice tails based on the length and nature of the lesion as shown in table 1. Value represent mean ± SD of 8 mice for *M. marinum* WT and 5 mice each for *H37Ra* and *M. smegmatis*. One way anova (***P=0.0001)

**Table 1:**
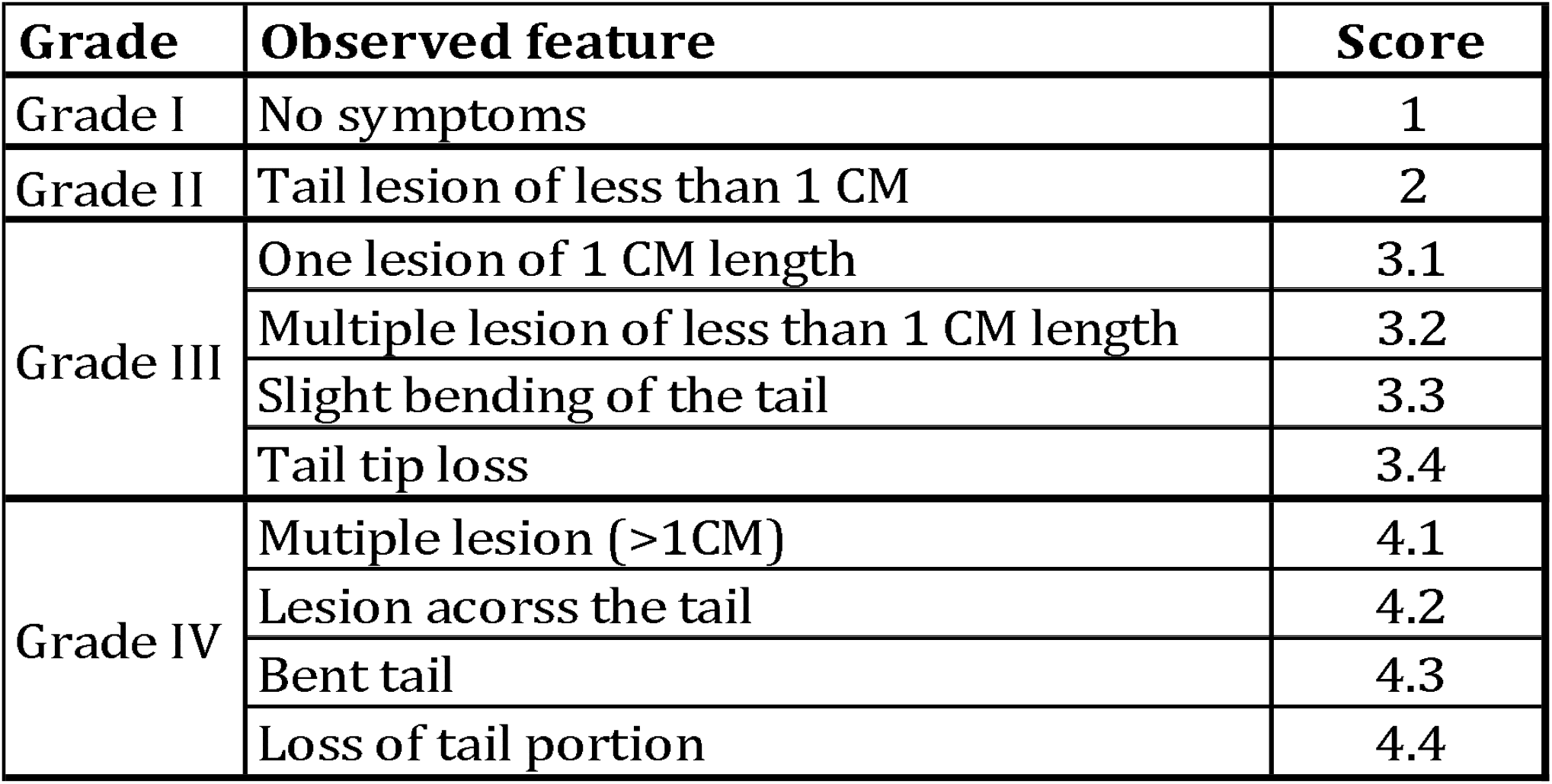
Grading of the observed pathological features.

| Grade | Observed feature | Score |
| --- | --- | --- |
| Grade I | No symptoms | 1 |
| Grade II | Tail lesion of less than 1 CM | 2 |
| Grade III | One lesion of 1 CM length | 3.1 |
|  | Multiple lesion of less than 1 CM length | 3.2 |
|  | Slight bending of the tail | 3.3 |
|  | Tail tip loss | 3.4 |
| Grade IV | Mutiple lesion (>1CM) | 4.1 |
|  | Lesion acorss the tail | 4.2 |
|  | Bent tail | 4.3 |
|  | Loss of tail portion | 4.4 |

### Tail lesions and bone erosion

The *M. marinum* infection to the mice tails is not limited to the soft tissue indeed it reached to the tail bone and caused erosion of the coccygeal vertebrae to the extent that the integrity of the some vertebrae is completely lost. The destruction of the tail vertebrae is not due to mechanical force since the erosion in the vertebrae is very much in line with the soft tissue damage (Fig. 2D) and also injection of PBS in mice tails has never resulted in any kind of damage. Whereas, the coccygeal vertebrae of the mice tails infected with *M. smegmatis mc*^*2*^ *155* and *H37Ra* exhibited typical bone architecture similar to that of uninfected mice tails (Fig. 2A) that suggests no bone erosion (Fig. 2B and 2C). The quantification of the bone erosion in terms of the bone mineral density (BMD) highlights the very significant drop in the BMD of mice infected with *M. marinum* while for *M. smegmatis mc*^*2*^ *155* the BMD drop is not significant

**Fig. 2:**
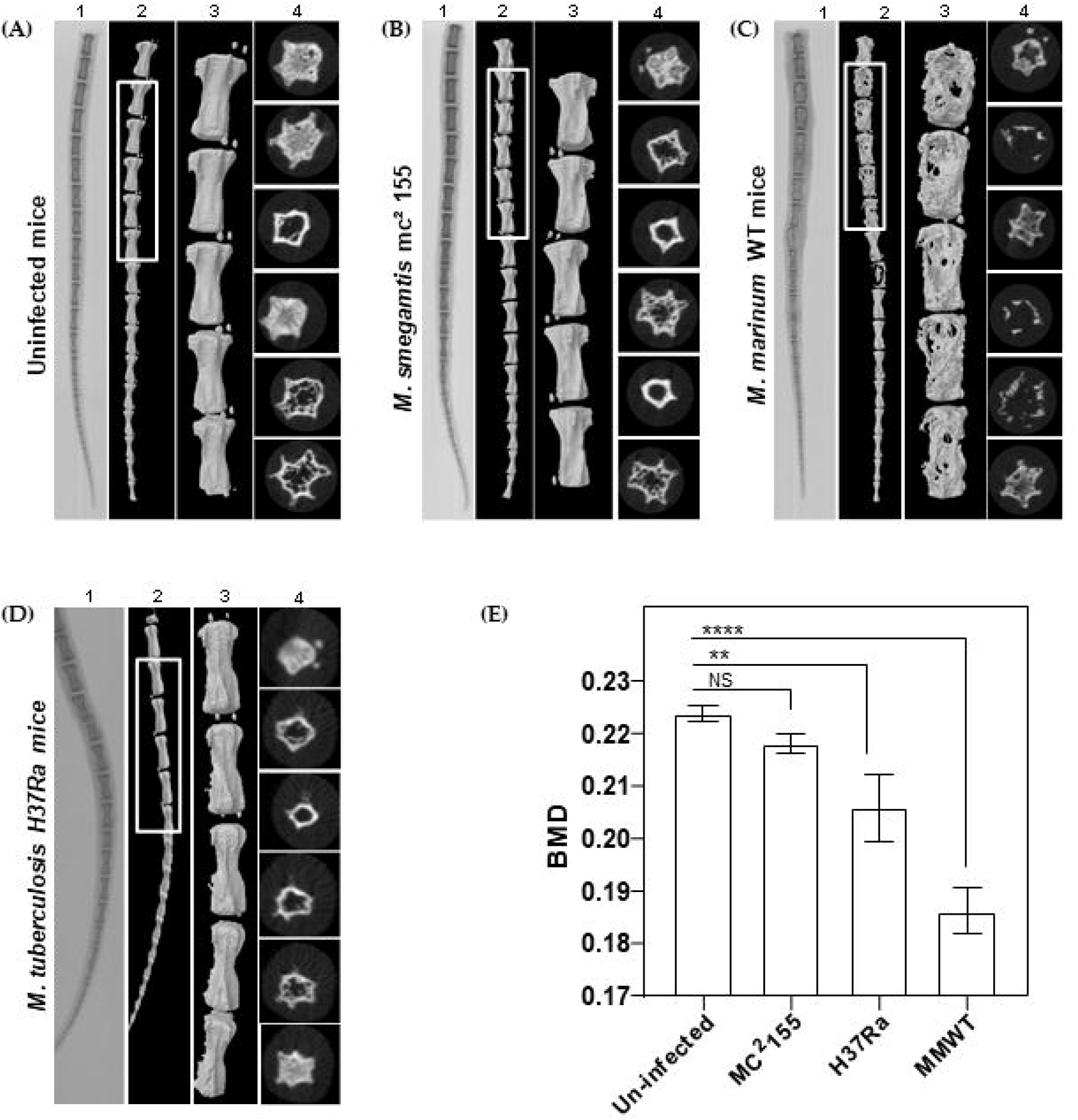
Micro-CT visualization of the infected mice tails. (A): Uninfected mice, (B): infected with *M. smegmatis mc*^*2*^ *155*, (C): *H37Ra* and (D): *M. marinum*. The mice tails were subjected to digital X-ray (panel labeled with 1) and Micro-CT (panel labeled with 2, 3 and 4 represents the whole mice tail, highlighted region of tail and representational images of the cross sections of the tail vertebrae respectively). (E): Quantification of the bone erosion in terms of bone mineral density (BMD) fo different *Mycobacterium species*. Values represent mean ± SD of 2 uninfected mice. While 3 mice for each *M. marinum* WT, *H37Ra* and *M. smegmatis* group. One way anova (****P<0.0001, P<0.01)

### Histological analysis of the tail lesions

The histopathological analysis of the sagittal sections of the mice tails using Hematoxylin and Eosin (H&E) staining revealed heavy infiltration of the immune cells and the loss of bony aperture in the *M. marinum* infected mice tails (Fig. 3C) that highlights the soft tissue damage and the bone erosion. The loss of bony aperture leads to the infiltration of the immune cells (mostly neutrophils) in the bone marrow space. The tail sections of the *M. smegmatis* and *H37Ra* infected mice exhibited the typical pathology similar to uninfected mice tails (Fig. 3A, 3B and 3D) as evident through the clear vertebral lining and the absence of the infiltration of the immune cell to the soft tissues and the bone marrow space. The soft tissue and the bone marrow space harbor the normal tissue architecture.

**Fig. 3:**
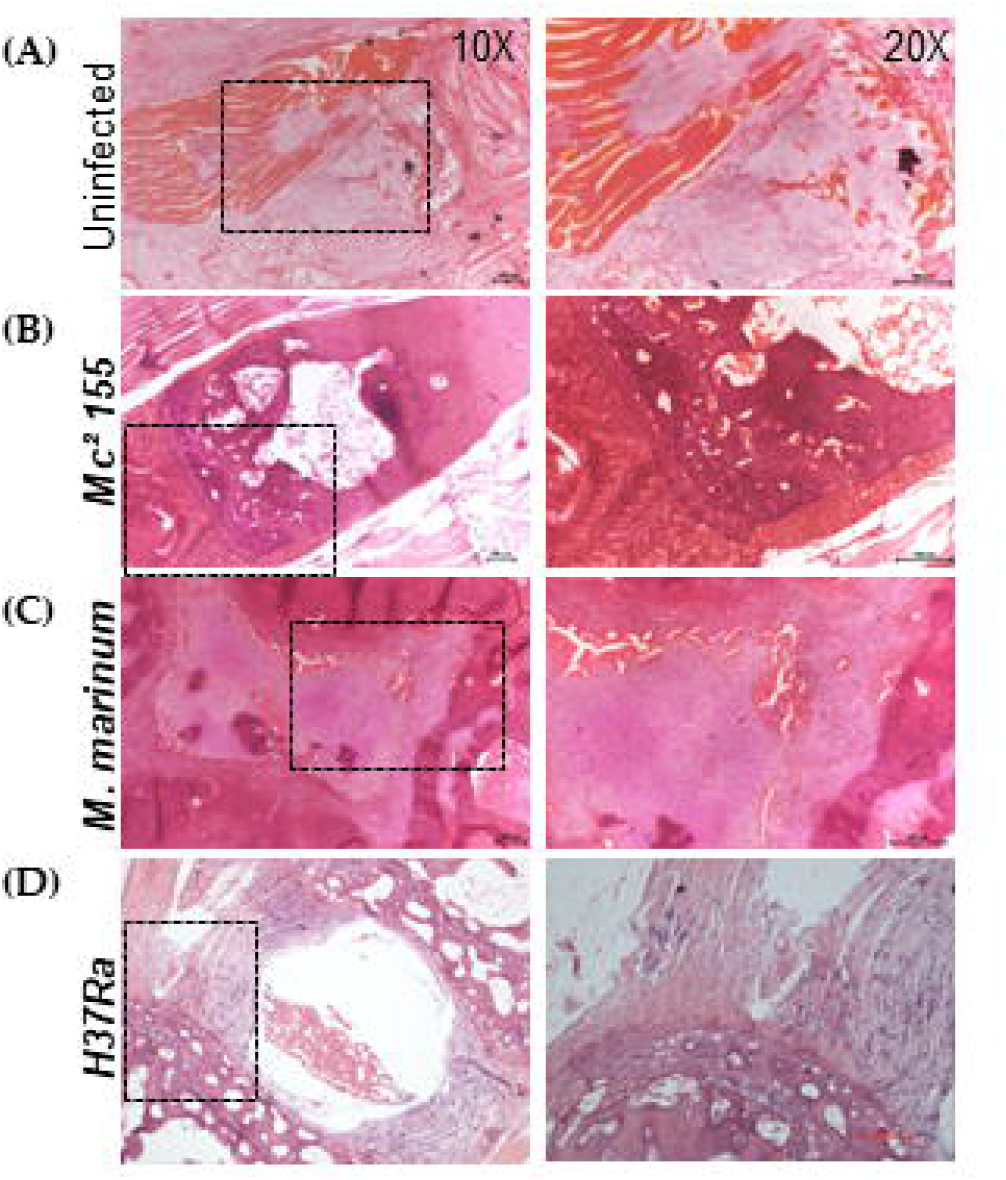
Histopathology of the infected mice tail sections: The sagittal section of the infected mice tail were obtained and stained with Hematoxylin and Eosin. (A): Uninfected mice tails sections (B): *M. smegmatis mc*^*2*^*155* infected mice tails sections (C): *M. marinum* infected mice tails sections.. (D): *H37Ra* infected mice tails sections. Where left panel shows the 10x image and the right panel the highlighted region at 20x.

### Histological analysis of lungs

The histopathological analysis of the sagittal sections of the mice lungs using Hematoxylin and eosin (H&E) staining revealed the formation of the granuloma like structure due to the infiltration of the immune cells in the *H37Ra* infected mice lungs (Fig. 4D). While, the *M. marinum* and *M. smegmatis* infected mice lungs possessed well-spaced alveolar space and the alveoli are intact and normal similar to the uninfected mice (Fig. 4A, 4B and 4C). The normal histopathology of the lungs of *M. marinum* and *M. smegmatis* infected mice suggests the nature and the site of the colonization of these mycobacterial strains, as *M. smegmatis* is non-pathogenic in nature and unable to colonize in any part of the mice body whereas, *M. marinum* is not able to survive in the lungs but it could successfully colonize and initiate strong inflammatory response in the tail where the temperature might be optimum for its growth. Akin to the above, the *H37Ra*, the attenuated counterpart of the laboratory strain *M. tuberculosis H37Rv*, is able to survive in the lungs as its optimum growth temperature is 37°C and evokes immune response for the formation of the granuloma like structure in the lungs as the alveolar spaces are filled with the immune cells, though the granulomatic structures are not atypical of the classical granuloma seen in the mice infected with virulent strains.

**Fig. 4:**
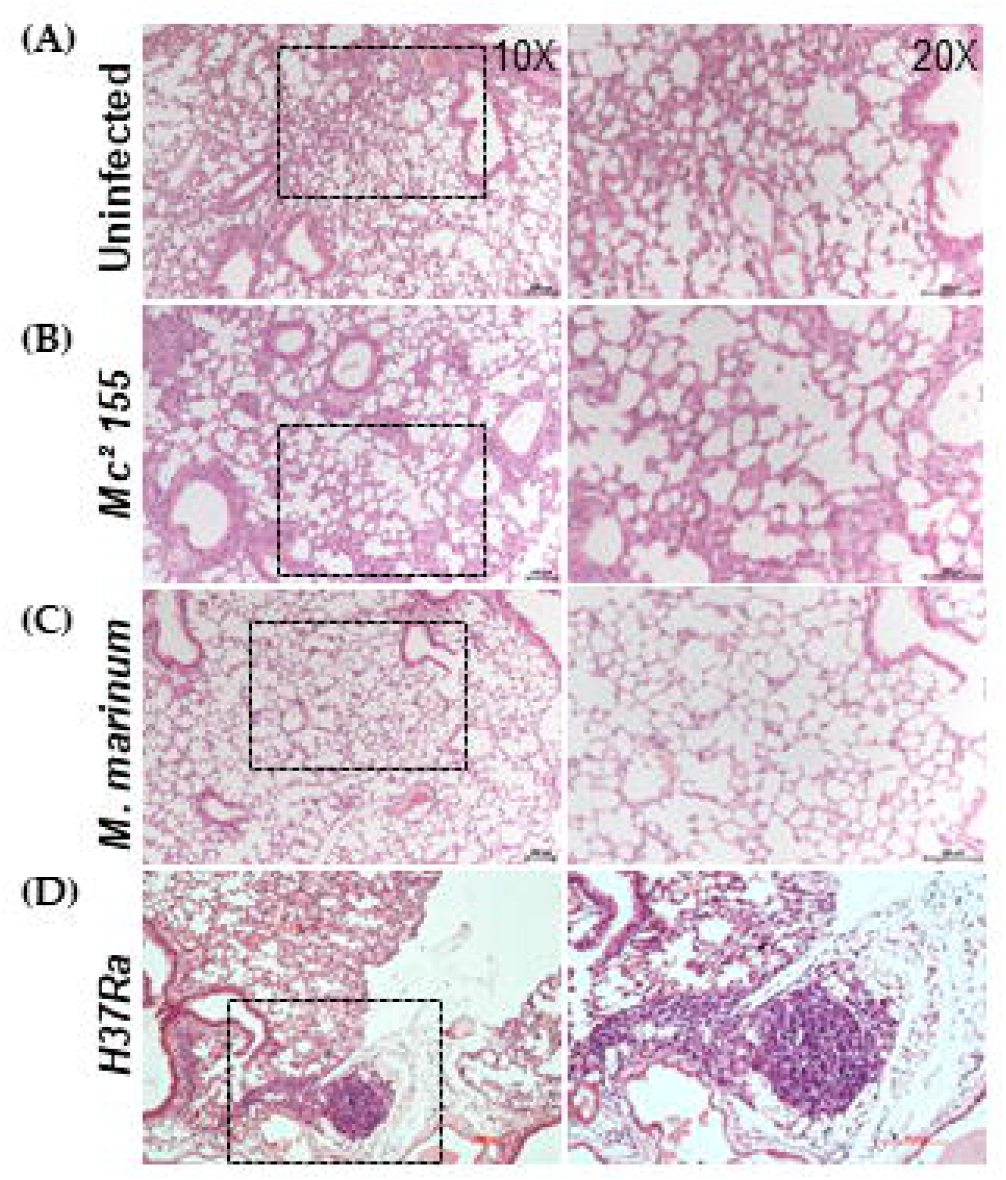
Histopathology of the infected mice lungs sections: The sagittal section of the infected mice lungs were obtained and stained with Hematoxylin and Eosin. (A): Uninfected mice, (B): *M. smegmatis mc*^*2*^*155*, (C): *M. marinum*, (D): *H37Ra*, Where, left panel shows the 10x image and the right panel the highlighted region at 20x.

### EmbRP Exhibited slow growth rate with extended lag phase

*M. marinum* WT was selectively grown in the presence of two different class of anti-mycobacterial drugs Ethambutol (EMB) and Isoniazid (INH) through a gradual increase in concentrations ranging from sub-lethal to lethal. The initial sub-lethal concentration was anticipated to allow the adaptation of the any pre-existing resistant form or the evolution of the new resistant form due to selection pressure of the drug. The culture thus selected at terminal concentration is termed as EmbRP and IRP for the antibiotics EMB and INH respectively. Furthermore, we also grew EmbRP in the absence the EMB to understand the withdrawal effect of the antibiotic on the EmbRP population; the population thus referred as EmbRP-Rev.

It can be seen from (Fig. 5A) that increase in concentration of EMB resulted in a gradual increase in lag time for the growth while a similar lag was not observed for IRP (Fig. 5B). We have also examined the growth of the ERP upon withdrawal of the EMB. It is clear from (Fig. 5C) that upon withdrawal of EMB, the growth of ERP is similar to that of the wild-type culture. We have also subjected the *M. marinum* WT through abrupt selection of the antibiotics at terminal concentration (8 μg/ml for EMB and 200 μg/ml for INH) and the *M. marinum* did not exhibit any growth till 25 days. The differences in the growth kinetics of the EmbRP and IRP may be due the differences in their mode of the action while EMB is bacteriostatic in nature, the INH is bactericidal. Thus, with the increase in the EMB concentration there is an increase in the lag phase that corroborates the time required for the adaptation of the bacterium in the presence of the antibiotic stress. However, upon withdrawal of the antibiotic, no significant change was observed in growth rate, in comparison to the *M. marinum* WT culture. It is pertinent to mention here that the *M. marinum* WT did not exhibit any growth when subjected to the terminal concentration indicating the importance of the adaptation.

**Fig. 5:**
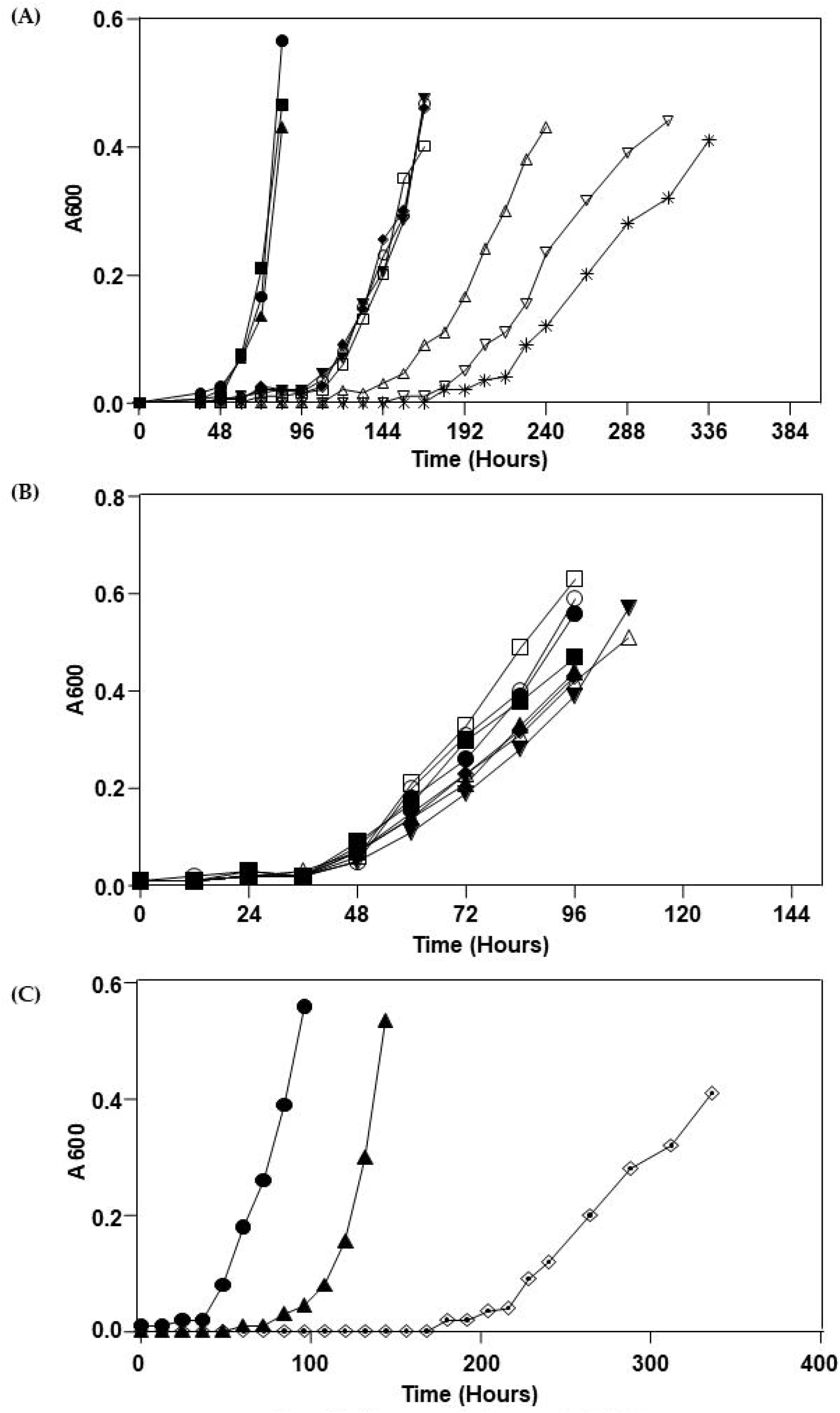
Growth curve of the anti-mycobacterial drug resistant *M. marinum* population. (A): *M. marinum* WT is cultured with gradual increase in the EMB concentration ranging from 0.00 μg/ml (●), 0.05 μg/ml (◼), 0.1 μg/ml (▴), 0.15 μg/ml (▾), 0.2 μg/ml (♦), 0.4 μg/ml (◯), 0.8 μg/ml (⊔), 2.0 μg/ml (△), 4.0 μg/ml (ν) and 8.0 μg/ml (*) to generate EMB resistant population (EmbRP). (B): *M. marinum* WT is cultured with gradual increase in the INH concentration ranging from 0.00 μg/ml (●), 2.0 μg/ml (◼), 4.0 μg/ml (▴), 8.0 μg/ml (▾), 20.0 μg/ml (♦), 50.0 μg/ml (◯), 100.0 μg/ml (⊔), 200.0 μg/ml (△) to generate INH resistant population (IRP). (C): Comparison of the growth kinetics of the different *M. marinum* population; *M. marinum* WT (●), EmbRP (◼) and EmbRP-Rev (▴).

### Infection of Mice with EmbRP and IRP

Infection of mice tails with EmbRP population resulted slow onset of infection and developed visible tail lesions of small size around 14 dpi (Fig. 6B) compared to mice infected with IRP and *M. marinum* WT that have rapid onset of symptoms emergence and the mice tail(s) developed lesions all across the tail length (Fig. 6A and 5C). The observed pathological features of the mice tails infected with IRP and EmbRP were graded based on the severity of the lesions as described earlier that suggests that there is no significant difference in the severity of the observed pathological features of the IRP infected mice; whereas the EmbRP infected mice showed significant reduction in the severity of the lesions compared with *M. marinum* WT infected mice (Fig. 6D).

**Fig. 6:**
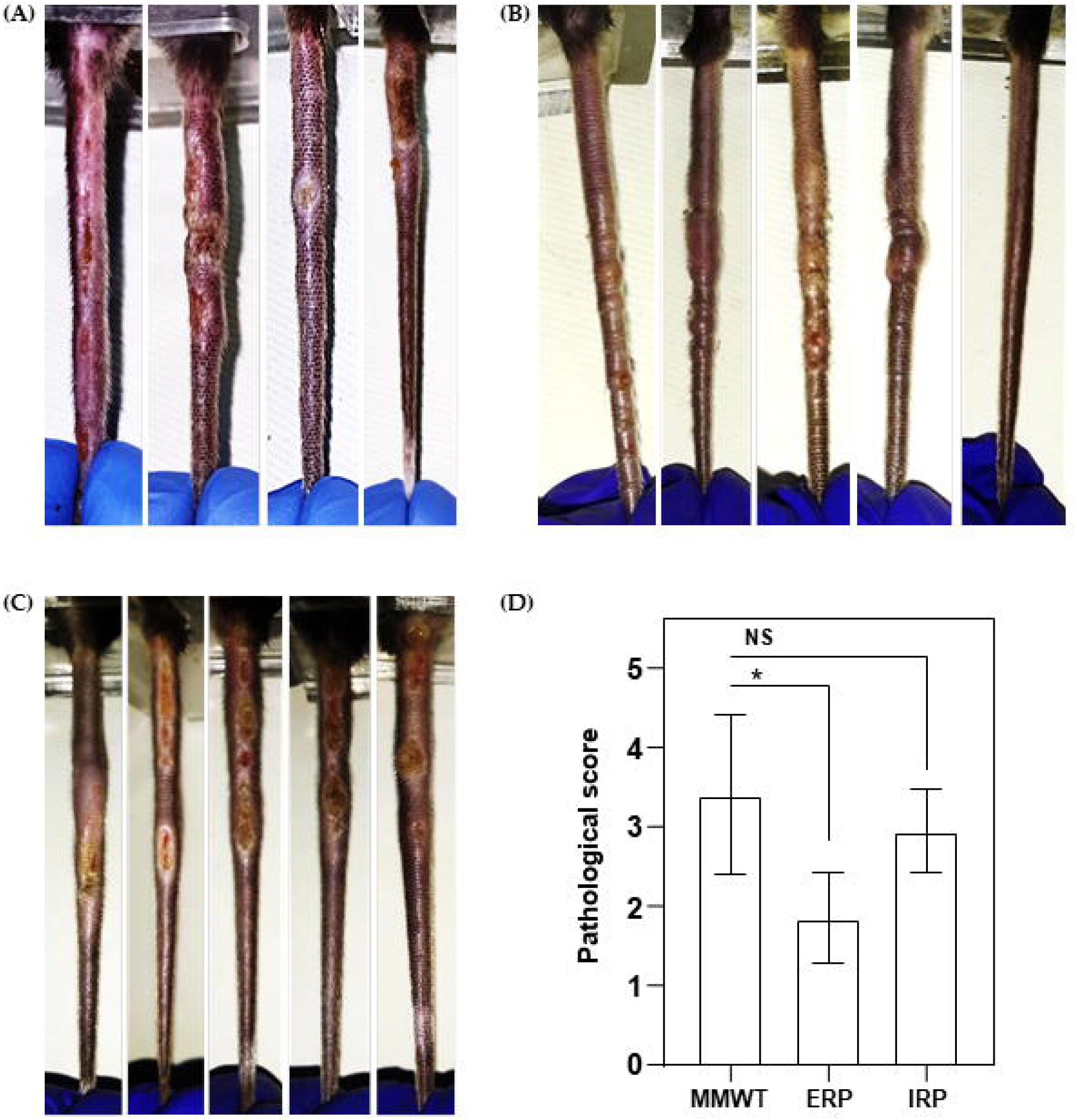
Mice infection with anti-mycobacterial drug resistant *M. marinum* population, via tail-vein injection with 1×10^7^ bacterial cells. (A): *M. marinum WT*, (B): *M. marinum* EMB resistant population (EmbRP), (C): *M. marinum* INH resistant population (IRP), and (D): quantitation of the severity of lesions of the infected tails based on the length and nature of the lesion as shown in Table 1. Values represent mean ± SD of 5 mice for each group *M. marinum* WT, EmbRP and IRP. One way anova (*P<0.05)

### Micro-CT to assess bone erosion of the mice tails infected with EmbRP and IRP

The ability to induce bone erosion of the EmbRP and IRP is further validated through micro-CT examination of the EmbRP and IRP infected mice tails. The mice infected with EmbRP did not show any significant degree of bone damage and bone erosion (Fig. 7C) and the vertebral architecture is intact like uninfected control mice tail (Fig. 7A). As oppose to EmbRP, the tails of IRP infected mice exhibited loss of vertebral architecture and significant bone erosion (Fig. 7D). Though, the nature of bone erosion is slightly different from the mice tails of *M. marinum* WT infected mice (Fig. 7B), as the bone become porous, it leads to a decrease in the bone content as, evident through the cross-sectional image of the vertebrae (Fig 7D.4) which subsequently resulted in the bone intensity below the normalization cutoff selected for the 3-D model preparation. The quantification of the bone erosion in terms of the bone mineral density (BMD) suggests a significant drop in the BMD of the IRP infected mice tails, while the drop in the BMD of EmbRP infected mice tails is insignificant (Fig. 7E). These findings suggest that the IRP is able to elicit the bone erosion similar to that of *M. marinum* WT; while, the EmbRP is not able to elicit to the same magnitude.

**Fig. 7:**
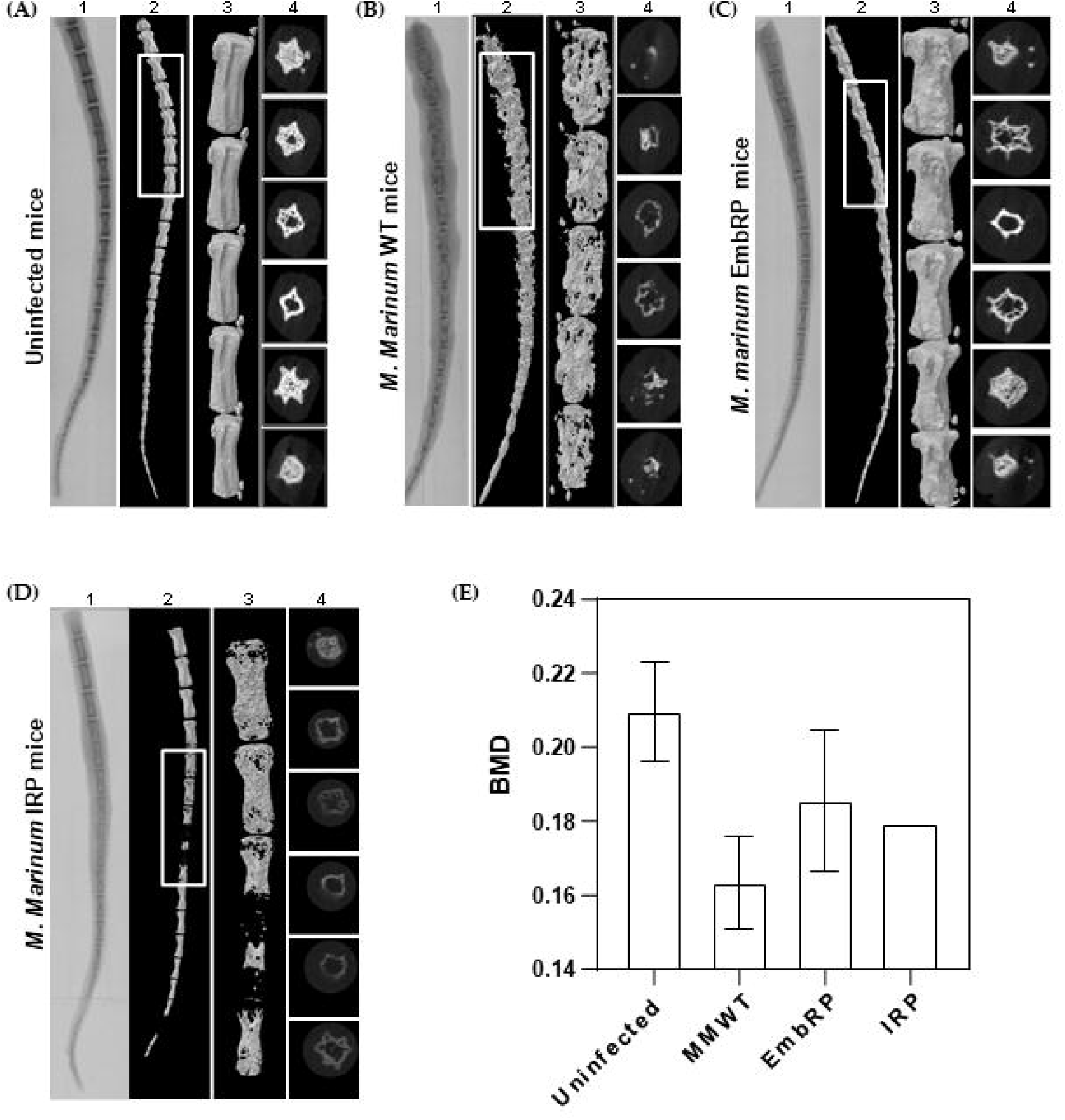
Micro-CT visualization of the mice tails of the mice infected anti-mycobacterial drug resistant *M. marinum* population. (A): Uninfected mice, (B): Infected with *M. marinum* WT, (C): *M. marinum* EMB resistant population (EmbRP), and (D): *M. marinum* INH resistant population (IRP). The mice tails were subjected to digital X-ray (panel labeled with 1) and Micro-CT (panel labeled with 2, 3 and 4 represents the whole mice tail, highlighted region of tail and representational images of the cross sections of the tail vertebrae respectively). (E): Quantification of the bone erosion in terms of bone mineral density (BMD) for different *M. marinum* population. Value represent mean ± SD of 2 mice for uninfected and *M. marinum* WT infected mice. While 3 mice for EmbRP and 1 mouse for IRP (P value is not significant).

### H&E staining of the section of mice tails infected with EmbRP and IRP

The mice tail sections of the EmbRP and IRP infected mice tails were also analyzed to assess the infiltration of the immune cells and the extent of the soft tissue damage. The sagittal sections of the EmbRP infected mice tails using Hematoxylin and Eosin (H&E) staining showed the infiltration of the immune cells to the soft tissues with the normal bone appendages like the uninfected control mice tail section (Fig. 8A) with no or minimal infiltration of the immune cells to the bone marrow space (Fig. 8C). Similarly, mice tail sections of the IRP infected mice tails showed infiltration of the immune cells to the soft tissues, with the erosion of the bone from the bone margin with no or minimal infiltration of the immune cells to the bone marrow space (Fig. 8D). The nature of the observed bone erosion is slightly different from the bone erosion observed in the mice tail section of the *M. marinum* WT infected mice tails that exhibited the loss of bone lining and infiltration of the immune cells to the bone marrow space (Fig. 8B).

**Fig. 8:**
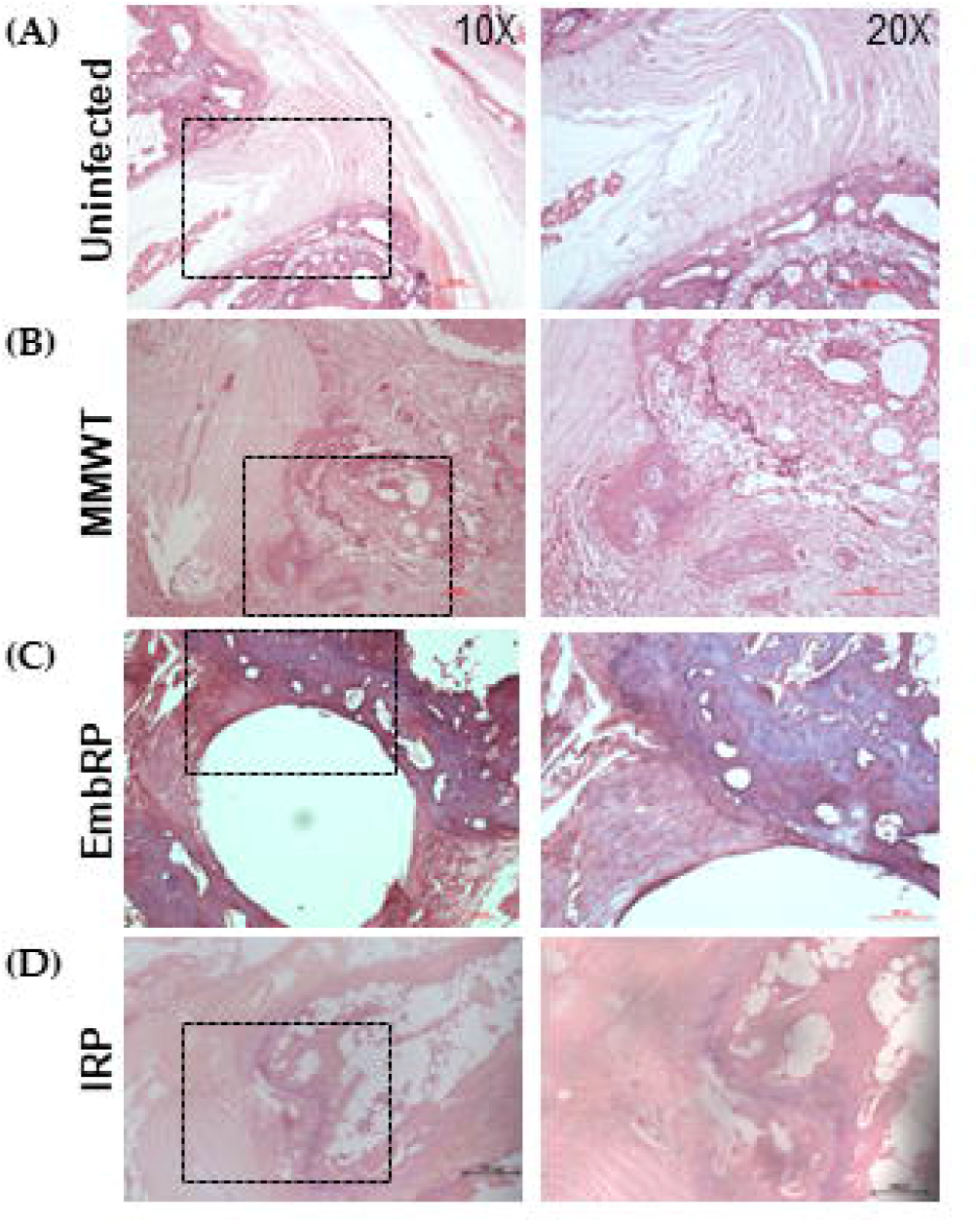
Histopathology of the infected mice tail sections: The sagittal section of the infected mice tail were obtained and stained with Hematoxylin and Eosin. (A): Uninfected mice tails sections, (B): *M. marinum* WT, (C): *M. marinum* EMB resistant population (EmbRP), (D): *M. marinum* INH resistant population (IRP). Where, left panel shows the 10x image and the right panel the highlighted region at 20x.

## Discussion

*M. tuberculosis* mainly colonizes inside the lungs and manifests the pulmonary TB but the infection can occur to other parts of the body to cause extra-pulmonary TB (EPTB). Several organs are affected by the EPTB that includes lymph nodes, gastrointestinal organs, bones and joints, central nervous system (CNS), and genitourinary organs [32,33]. Approximately 15-20% of the tuberculosis infections are EPTB [34,35]. The treatment and diagnosis of the EPTB is a challenging task as the classical acid fast mycobacterial staining or the mycobacterial culture techniques often returns negative in a laboratory setup [36]. The diagnostics further require more advanced approaches like nucleic acid test (NAT) tests, computed tomography scan (CT scan) or magnetic resonance imaging (MRI), though, it can be diagnosed by Tuberculin skin test (TST) and IFN-γ releasing assay (IGRA) but these methods offers very poor sensitivity which require further confirmation [37]. The diagnostic challenge is further exacerbated due to inaccessibility of the tissues that often requires invasive procedures for sample collection and often the radiological features of the symptoms resemble with other diseases leading to inaccurate conclusion [36,37]. The prominent association of paradoxical reaction to EPTB than pulmonary TB poses more difficult challenge to decide most accurate therapeutic regimen [35,37,38]. Bone and joint TB/skeletal TB is one such EPTB which is characterized by the colonization of the bacterium to the bony tissues mostly spine but it can infect the whole skeleton like hip joint, knee joint etc. [39]. Mostly, infection initiates from the trabecular bone of the epiphysis but sometime it may originate from the central or anterior part of the vertebrae [35]. The progression of the disease is characterized by the destruction of the epiphyseal cortex, intervertebral disc and spread to adjacent vertebrae. Sometimes, in severe cases of infection, the whole vertebral shaft gets eroded and become soft leading to the loss of vertebrae [35].

The presence of a reliable and reproducible *in vivo* model for EPTB, especially bone is of utmost importance to understand the onset and progression that resembles the scenario in human host [22,31]. The first use of *M. marinum* to infect mouse by shepherd’s group in 1963 has demonstrated the use of this model to understand the lesion development and its possible use in understanding the tuberculosis pathogenesis [40,41]. Another study reported by the Eric Brown and colleagues [31] described the kinetics and onset of the granuloma formation together with the loss of coccygeal vertebral bone volume of the infected mice tails [31,42,43]. The reliability of *M. marinum* infection of mice deserves a closer examination in view of the interesting outcome of the pathology such as bone erosion through associated histopathological observations. The present study aimed to understand the reliability of this approach regarding the onset and progression of the extra-pulmonary infection (in this case bone TB) with the cognizance of the histological analysis to demonstrate the nature and extent of the bone erosion and underlying populations while a treatment regimen is in place.

The data presented in the Figures 1 to 4 depict the characterization and manifestation of the pathological features with the onset of the infection in general but bone-damage *per se*. Among the well-studied mycobacterial strains such as *M. marinum, H37Ra* and *M. smegmatis mc*^*2*^ *155*, only infection of mice with *M. Marinum* has elicited extensive lesion formation due to soft-tissue damage as well as erosion of bones to extent of disappearance [31,43]. The H&E staining of the several tail sections of the numerous infected mice tails has revealed dramatic infiltration of the immune cells to the bone marrow space together with the soft tissue with a gradual to rapid erosion of the coccygeal vertebrae, unambiguously confirmed by the micro-CT imaging of numerous tails. This approach always showed significant bone erosion of the mice infected with *M. marinum* while mice infected with *H37Ra* and *M. smegmatis mc*^*2*^ *155* displayed normal bony aperture as well as no infiltration of immune cells. For example, the histopathological examination of lungs of mice infected with *M. marinum* and *M. smegmatis mc*^*2*^ *155* revealed the normal tissue architecture with distinct alveolar spaces, however, infiltration of the immune cells is observed in the sections of the *H37Ra* infected mice lungs that resulted in the formation of granuloma like structure [44]. These findings further suggest that *M. marinum* grows/adopts inside the cooler part of the mice and are unable to form the niche inside the body. Whereas, the *H37Ra* despite being the attenuated counterpart of *M. tuberculosis H37Rv*, it forms granuloma like structure in lungs [44–46].

Among many possibilities that can contribute to the evidenced pathologies orchestrated by the *M. marinum*, the possibility of *M. marinum* having micro/sub populations that have the ability to colonize efficiently in mice tails better than other organs exists in principle. It is therefore, we wished to examine whether or not treatment with INH or EMB, used extensively as the first line of drugs against tuberculosis infection, leaves or retains some bacterial population that can still inflict soft-tissue and/or bone-damage during recovery period in which no bacterial treatment regimen will be in place.

A clear answer to this question is essential as elimination of mycobacteria after the first round of treatment with these drugs can still leave a small/residual bacterial population/colony that can cause soft-tissue/bone-damage or even re-emerge as a resistant variety [47,48]. We, therefore, further assessed the infectivity potential of the antibiotic resistant-population generated *in vitro*. We selectively cultured *M. marinum* in the presence of selection pressure of the two frontline anti-mycobacterial drugs INH and EMB through gradual increase of the concentration ranging from sub-lethal to lethal concentration which is above its known MIC_90_. The *M. marinum* forms, thus selected, were further used to infect the mice and assessed for their virulence potential as well as their capacity to keep virulence determinant intact over the antibiotic selection. The data depicted in Figures 5-8 represent the growth kinetics *M. marinum* selected populations (EmbRP and IRP) and their infection to the mice tails. The pathological features of the mice tails infected with IRP are very much similar to that of the infection with *M. marinum* WT, whereas, the EmbRP infected mice showed significantly delayed onset of the symptoms. Furthermore, the mice infected with IRP exhibited bone erosion at the same time the bone lining seems to be intact with normal bone marrow space but on a similar scale the EmbRP infection could not result in significant bone erosion as compared to *M. marinum* WT with intact vertebral lining and periosteum, together with the well-spaced bone marrow with less infiltration of the polymorphonuclear cells together with other immune cells. These findings suggest that IRP has retained the virulent determinants of *M. marinum* WT whereas the EmbRP population might have lost some virulent determinant(s) responsible for the lesser tissue damage/bone-erosion. While it can be argued that the delayed onset of symptoms by EmbRP could be due to its slow *in-vitro* growth. We rule out this possibility because the EmbRP upon withdrawal of the drug did not exhibit any lag in its growth in comparison to the wild-type *M. marinum* and also the mice were not administered EMB through any route.

*M. marinum* infection to mice resulted distinct pathological features not observed in the murine tuberculosis model with extensive bone erosion, it is expected to open a new approach to understand the progression of extra pulmonary TB, precisely colonization and/or erosion of bone by the tuberculosis bacilli. This approach can be further utilized to screen small molecule(s) or therapeutic agents that can yield information required to develop new therapeutic approaches [31,43]. This approach can further strengthen the available model systems to decipher deeper information on the bone TB [49–51]. There are incidences of mice infection with other *Mycobacterium* spp. like *M. tuberculosis and M. ulcerans* causing osteomyelitis. Moreover *M. tuberculosis* infected mice don’t exhibit typical caseating granuloma together with the lack of distinct histological features as observed in the presented model, whereas, *M. ulcerans* infected mice exhibited, slow growth rate and delay in the emergence of symptoms requiring longer investigation time as compared to present study [52–54]. Although, the zebrafish infection with *M. marinum* exhibits spinal curvature [55,56] the underlying causes are still not clear. *M. marinum* infection to mice offers visible and quantifiable pathological features in the form of tail lesions as early as 4-7 dpi and culminate around 15-18 dpi. Furthermore, the micro-CT and histopathological images confers the loss of bony aperture and infiltration of the immune cells to form granulomatous structure; importantly, the histological sections revealed the loss of periosteum and infiltration of the immune cells to the bone marrow space.

In summary, the *M. marinum* infection of mice tails is highly informative regarding the bone erosion and probably the factors that lead or accelerate the onset of disease symptoms. The manifested symptoms are akin to skin lesion or the osteomyelitis of the individuals infected with *M. marinum* that can be employed to understand the progression of *M. marinum* infection to humans.

## Acknowledgements

The authors thank Dr. B. Ramana Murthy for assistance in *in vivo* experiments, Dr. Mohan Wani and Mr. Shubhanath Behera for Micro-CT visualization and National Centre for Cell Science and Department of Biotechnology for generous intra-mural support. MK, RA, PU are recipients of senior research fellow ships of CSIR, India.

